# Identification of Beneficial Bacterial Strains for Tomato Growth Promotion and Biocontrol of Bacterial Canker Caused by *Clavibacter michiganensis*

**DOI:** 10.64898/2026.06.01.729334

**Authors:** Nasim Sedighian, Marie-Christine Groleau, Eric Déziel

**Affiliations:** Centre Armand-Frappier Santé Biotechnologie, Institut National de la Recherche Scientifique (INRS), Laval, Québec, Canada

**Keywords:** biological control, sustainable agriculture, bacterial canker disease, plant growth–promoting bacteria (PGPR), secondary metabolites, phylogenomic

## Abstract

Bacterial canker of tomato, caused by *Clavibacter michiganensis* (*Cm*), remains difficult to control due to lack of effective management options. In this study, a collection of over 500 bacterial isolates was screened *in vitro* for antagonistic activity against *Cm* and plant growth–promoting (PGP) traits. Based on these results, 32 candidates were evaluated *in planta*, leading to the identification of three highly effective strains: *Pantoea agglomerans* SO16PY and two *Pseudomonas marginalis* sensu lato strains, IRDA16 and SO16PC, which consistently enhanced tomato vegetative growth. Notably, *P. agglomerans* SO16PY delayed disease onset in *Cm*-inoculated plants by up to 7 days and significantly reduced wilting severity, lowering the disease severity score from 85% to 45%. Strains IRDA16 and SO16PC also restricted disease development, reducing severity scores to 67.5% and 57.5%, respectively. Whole-genome sequencing and comparative genomics revealed that strains IRDA16 and SO16PC form a distinct, specialized rhizosphere lineage within the *Pseudomonas marginalis* group, exhibiting average nucleotide identity (ANI ≈ 96%) and digital DNA-DNA hybridization (dDDH ≈ 69.5%) values near species delineation thresholds. Genome mining identified diverse biosynthetic gene clusters (BGCs) encoding non-ribosomal peptide synthetases (NRPS), the lipopeptide viscosin, and terpenes, which likely drive the biostimulant and antagonistic traits of this novel *Pseudomonas* lineage. Together, these findings characterize promising bacterial candidates with dual biostimulant and biocontrol capacities while uncovering a genomically distinct *Pseudomonas* lineage optimized for beneficial plant–microbe interactions in sustainable agriculture.

**IMPORTANCE:** *Clavibacter michiganensis* (*Cm*) is a major bacterial pathogen of tomato and poses a significant economic threat to global production. It is classified as an A2 quarantine pathogen by the European and Mediterranean Plant Protection Organization (EPPO). Current management strategies rely largely on chemical control, including copper-based compounds (e.g., Bordeaux mixture, copper oxychloride), mancozeb, and antibiotics like streptomycin. However, these approaches raise increasing concerns related to environmental contamination, phytotoxicity, and the development of resistant pathogen populations. As a sustainable alternative, plant growth–promoting bacteria (PGPR) have emerged as promising biocontrol agents. In this study, we identified bacterial strains exhibiting antagonistic activity against *Cm* both *in vitro* and *in planta*. Notably, these strains also enhanced tomato growth parameters, demonstrating their dual functionality. Given the environmental drawbacks associated with chemical inputs, the use of such beneficial microorganisms represents a promising strategy for advancing sustainable and ecofriendly tomato production systems.

---

Tomato (*Solanum lycopersicum*) is the second-most consumed vegetable crop in the world after potato, with an annual production of 192 million tons in 2023 (1).

Over the last decades, demands have been growing toward green agriculture and using healthier products, free of synthetic agrochemicals (2). Therefore, an important goal in modern agriculture is to develop more effective and safer alternatives to fight plant diseases, while still improving crop quality and production (2). Within this frame, biocontrol represents an attractive ecofriendly and a sustainable avenue to control phytopathogens. In this approach, organisms or products from biological origin are used, such as natural enemies or biobased products derived from microbes, plants or animals, to control pests (3). A substantial proportion of biological control agents belong to the group of plant growth-promoting rhizobacteria (PGPR). They can improve the general plant health status through a variety of mechanisms, including phosphate and potassium solubilization, indole-3-acetic acid (IAA) production, nitrogen fixation, siderophore production and aminocyclopropane-1-carboxylate (ACC) deaminase activity (4,5). In addition, they colonize plant roots and exclude phytopathogens through means such as competition, induction of resistance, siderophore production, and secretion of secondary metabolites, hydrolytic enzymes, and volatile compounds (6,7). Another advantage to this approach is that resistance development to biopesticides is less likely because they have complex, often multiple, modes of action (competition, parasitism or production of secondary metabolites) that pests cannot easily overcome (8,9,10).

Bacterial canker of tomato, caused by the Gram-positive bacterium *Clavibacter michiganensis* (*Cm*), is considered one of the most devastating plant diseases worldwide. This phytopathogen is listed on the A2 list of quarantine pathogens (11). The disease caused by *Cm* was first described in 1910 on tomato plants in Michigan (USA) (12) and is currently widespread worldwide (11,13). Bacterial canker is responsible for significant economic losses for tomato growers (14,15,16). Diverse approaches are used to control this disease but until now, only one product is commercialized as a dedicated bioprotectant (Agriphage CMM™) (3).

To date, there is no report regarding beneficial bacterial strains isolated in North America against bacterial canker. Moreover, functionality of each PGPR may vary in different climate conditions and locally adapted strains yield better field performance. Therefore, in this study, we analyzed different bacterial strains, isolated in North America (Canada and U.S.) to (i) assess their PGP and *Cm* biocontrol potential *in vitro*; (ii) evaluate their effect on tomato growth; (iii) control the development of bacterial canker disease on tomato plants; (iv) acquire the complete genome sequence of promising isolates to identify biosynthetic gene clusters and other genes that could be involved in biocontrol and biofertilization activities. This dual screening enabled the identification and initial characterization of multifunctional strains with biofertilization and biocontrol potential.

## RESULTS

### Identification and *in vitro* characterization of bacterial strains

In order to identify bacterial strains with both antagonistic activity against *Cm* and plant growth-promoting (PGP) traits, more than 500 candidates were tested *in vitro*. In the initial screening, approximately 250 isolates showed a measurable antagonistic activity, as indicated by the formation of inhibition zones against a *Cm* lawn on agar plates, ranging from 103.8 to 881.4 mm^2^.

These isolates were further assessed for their PGP potential. Among those, we found 32 candidates who also expressed PGP functions *in vitro* (Table S1). Sequencing of the gene encoding for the 16S rRNA showed that they belonged to the following genera: *Pseudomonas* (65% of candidate strains), *Burkholderia* (9%), *Pantoea* (12%), *Bacillus* (9%), and *Erwinia* (3%). One *Pantoea* isolate (SO16PY) displayed the largest (881.4 mm^2^) inhibition zone against *Cm* growth (Table S1).

### Evaluation of PGP activities of bacterial strains *in planta*

Following primary *in vitro* screening, 32 promising bacterial candidates were selected for further evaluation of their plant growth-promoting effects *in planta*. Two tomato cultivars, Trust and Heinz 2206, were cultivated in a growth chamber using two different soil conditions (100% PROMIX and 70% PROMIX : 30% sand). In the first screening using cv. Trust (PROMIX), strains *Pseudomonas donghuensis* M3, *Bacillus pumilus* 615, *Pseudomonas orientalis* 9, *Pantoea agglomerans* SO16PY, and two *Pseudomonas marginalis* sensu lato isolates (IRDA16 and SO16PC) showed significant positive effect on tomato vegetative growth (Table S2). However, following repeated experiments, only two isolates of *P. marginalis* sensu lato, strain IRDA16 (isolated from apple leaf, 2015, Quebec) and strain SO16PC (isolated from tomato leaf, 2016, Quebec), as well as *P. agglomerans* strain SO16PY (isolated from tomato leaf, 2016, Quebec) significantly and consistently promoted tomato root and shoot parameters *in planta* (Fig. 1; one repetition is shown in supplemental figure S1). For cultivar Trust using PROMIX, both strains of *P. marginalis* significantly improved fresh/dry weight of root and shoot. On the other hand, *P. agglomerans* SO16PY was especially efficient at increasing fresh/dry weight of root and shoot of both cultivars with soil diluted with sand, while in PROMIX, it only significantly improved fresh weight of root and shoot of cultivar Heinz 2206 (Fig. 1). *P. marginalis* SO16PC also improved fresh/dry shoot weight under less rich soil condition for both cultivars, indicating its adaptable potential under environmentally different soil conditions. In PROMIX, strain IRDA16 increased fresh and dry root/shoot weight in cultivar Heinz 2206.

**Figure 1.**
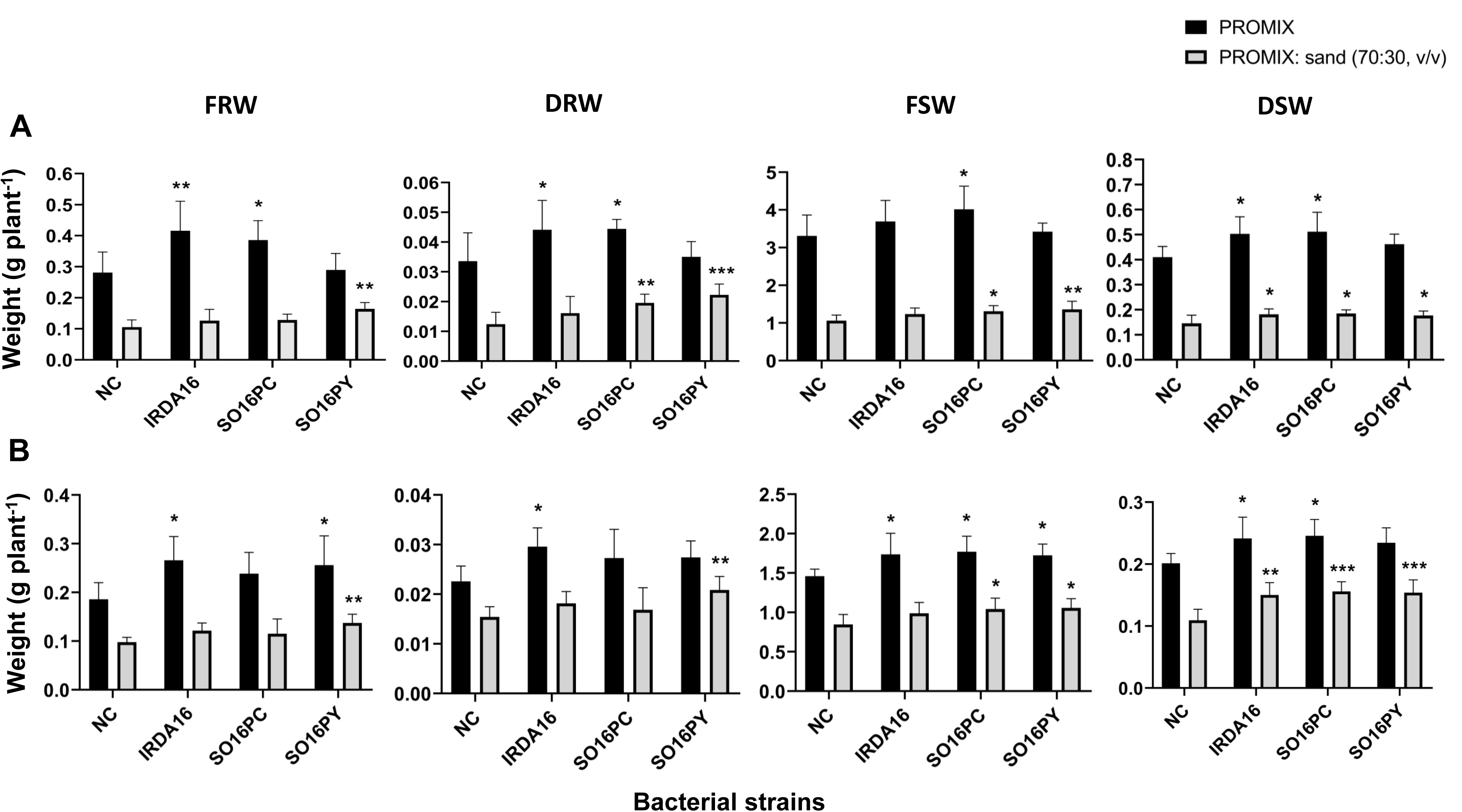
Plant-growth promoting effect of bacterial candidates on tomato growth parameters. **A.** cv. Trust, and **B.** cv. Heinz 2206. Axis X and Y respectively show bacterial strains and plant organs weight per gram. FRW: fresh root weight, DRW: dry root weight, FSW: fresh shoot weight, and DSW: dry shoot weight. Bars indicate means ± standard errors (n=7). NC, negative control (non-inoculated plants treated with sterile distilled water). Data presented is a combination of two independent repeats. Asterisks show the significant difference compared to NC: *, p ≤ 0.05; **, p ≤ 0.01; and ***, p ≤ 0.001.

Strains IRDA16, SO16PC and SO16PY also exhibited strong antagonism against *Cm,* with inhibition zones of 572.5, 551.5 and 881.4 mm^2^, respectively (Table S1 and Table 1). They also exhibit several key PGP traits including siderophore production, phosphate and potassium solubilization and ACC deaminase activity (Table 1). In addition, strain SO16PY produced a high level of IAA (23.7±0.4 µg/ml) in liquid LB medium supplemented with 5 mM L-tryptophan. On the basis of the combined *in vitro* (Table 1) and *in planta* (Fig. 1) results, these three strains were selected for further evaluation in subsequent experiments.

**TABLE 1.**
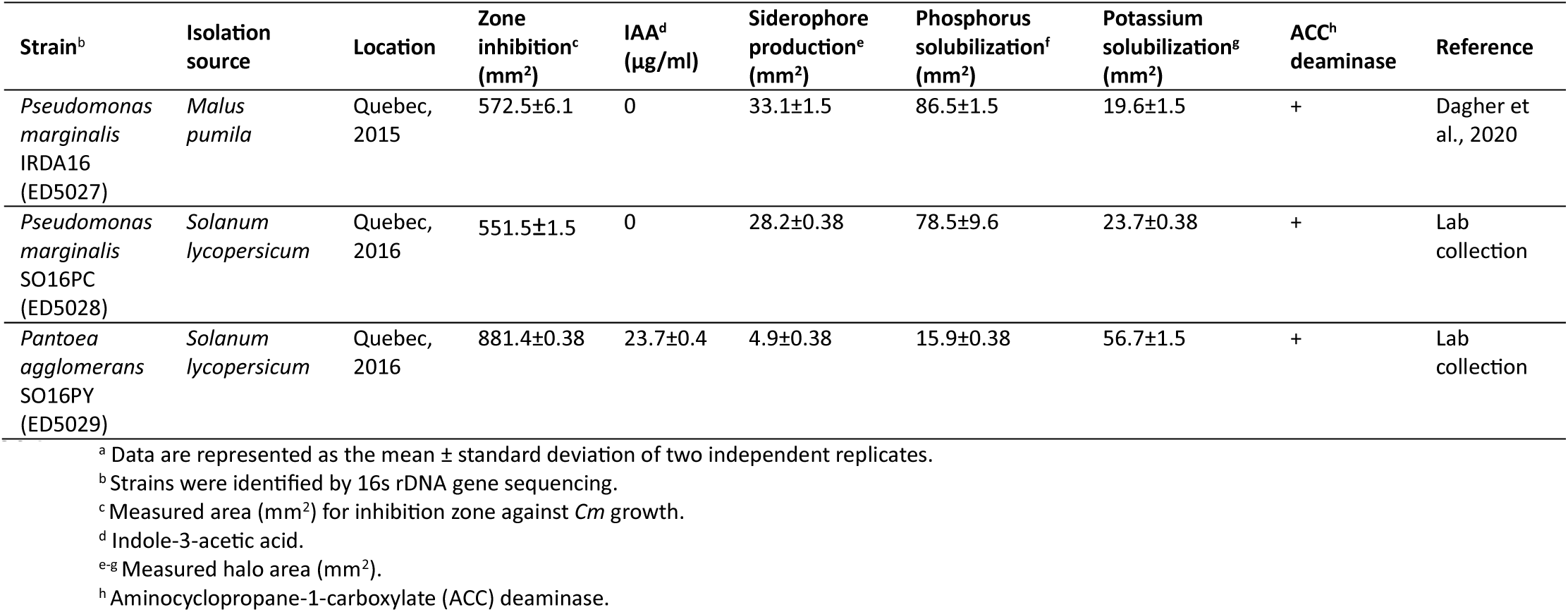
Plant-growth promoting characteristics of selected bacterial strains *in vitro*.

### Bacterial candidates improve tomato primary root length *in vitro*

Strains IRDA16, SO16PC and SO16PY were further used to assess their effects on tomato seed germination. All three bacteria had no negative impact on seed germination (cultivar Trust), and even significantly improved primary root length *in vitro* (Fig. 2).

**Figure 2.**
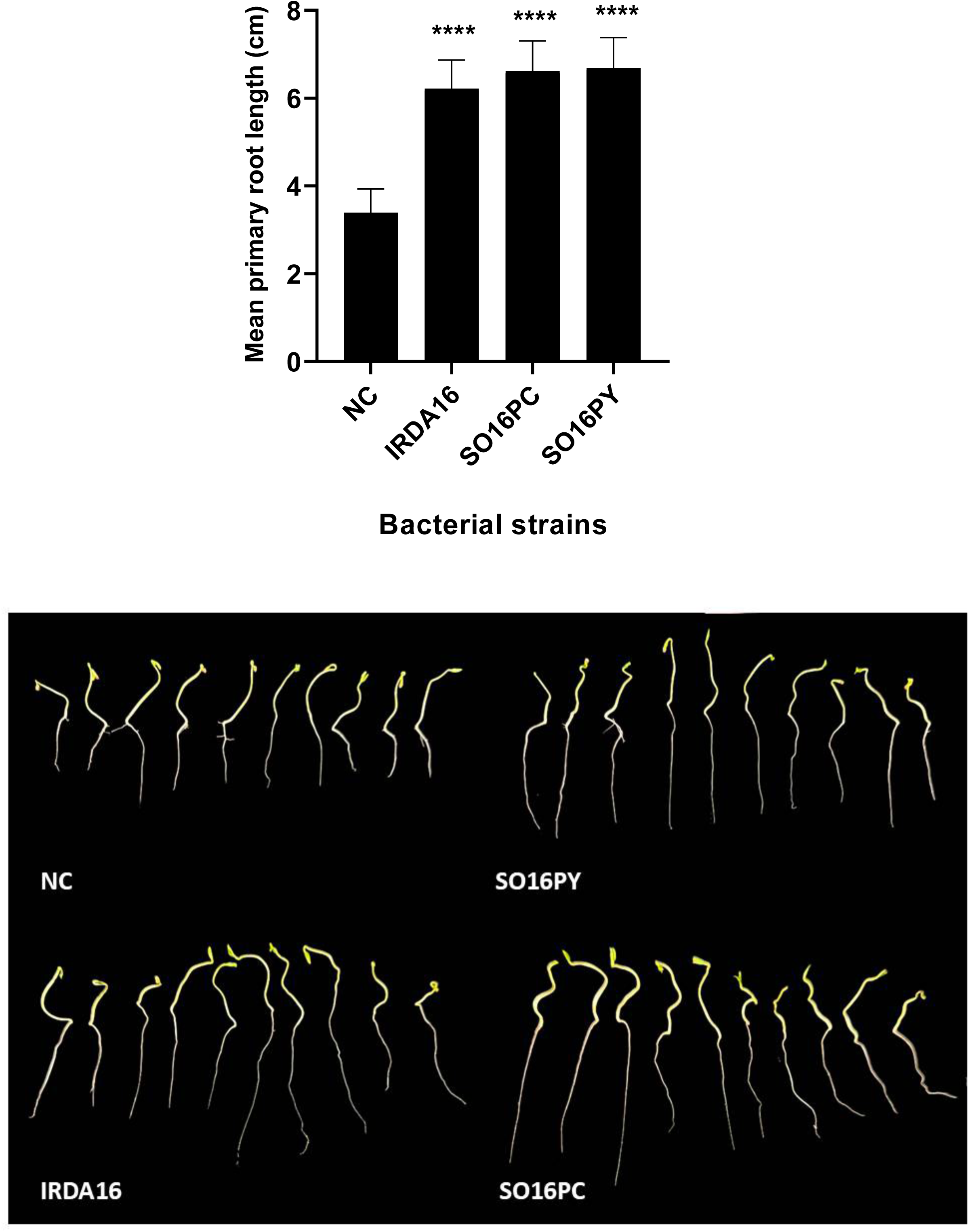
**Plant-growth promoting effect of bacterial candidates in tomato seed germination**. Bars indicate means ± standard errors (n=20), from two replicates. NC, negative control (non-inoculated seeds treated with sterile distilled water). Asterisks show the significant difference compared to NC: ****, p ≤ 0.0001.

### Antagonistic effect of cell-free supernatants

The clear inhibition zones produced by the three bacteria against *Cm* grown on TSA plates (Table 1) suggested the release of diffusible antimicrobial compounds. To verify the presence of extracellular inhibitory metabolites*, Cm* liquid cultures were supplemented with cell-free spent supernatants of the three selected bacterial strains (IRDA16, SO16PC and SO16PY). All three supernatants completely inhibited the growth of *Cm* strain LMG7333^T^ (Fig. 3). In contrast, spent supernatant of the negative control *E. coli* DH5α had no effect on growth of *Cm*.

**Figure 3.**
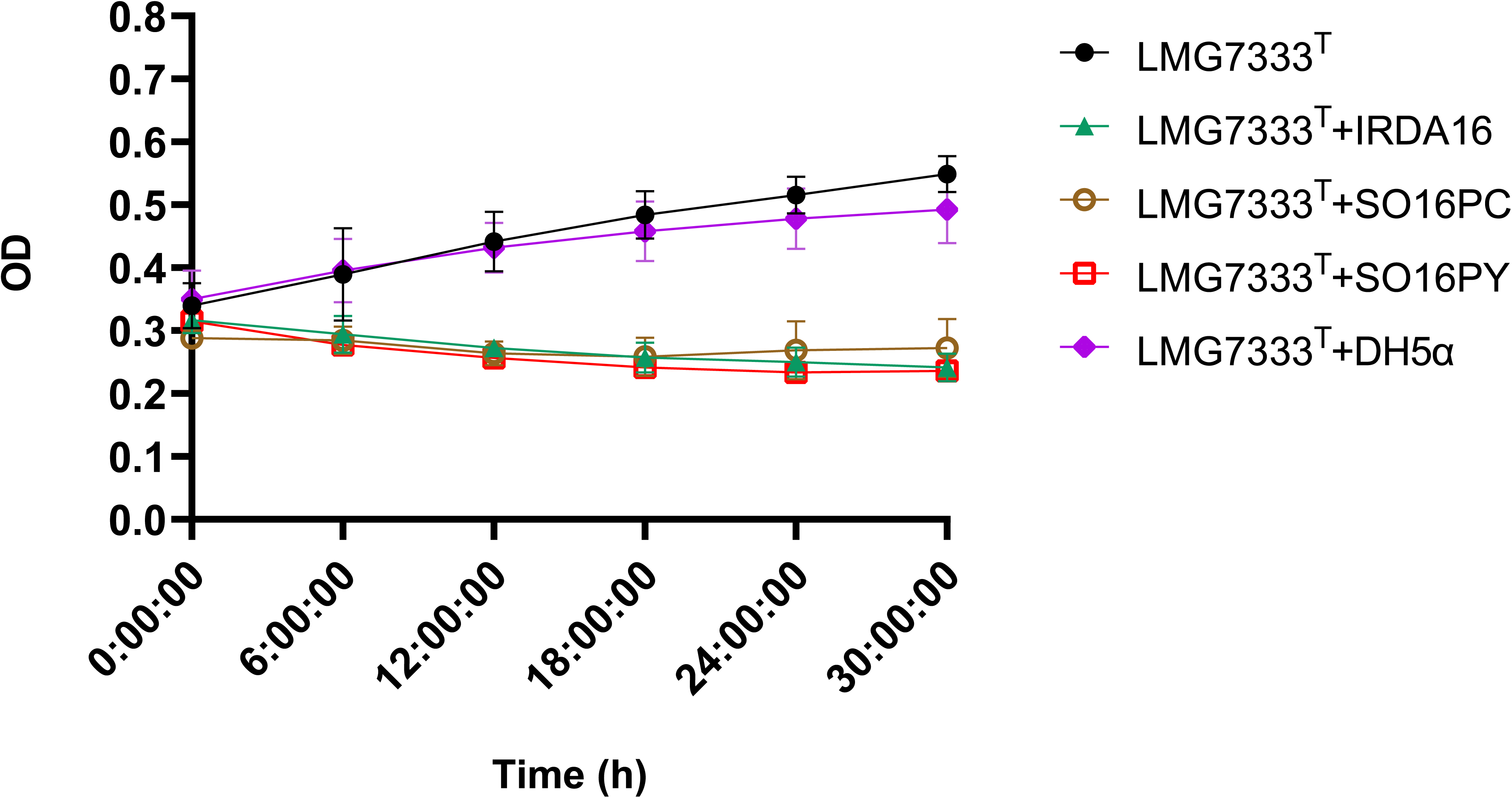
Inhibitory effect of cell-free spent supernatants of bacterial strains against *Cm* growth *in vitro*. Bars indicate means ± standard errors (n=6), from two replicates. Optical densities (OD_600_) were automatically measured by Bioscreen every 6 hours. *Escherichia coli* DH5α was used as negative control.

### Evaluation of biocontrol activity of bacterial candidates against *Cm in planta*

Following confirmation of antagonistic activity against *Cm* growth *in vitro* and expression of plant growth-promoting traits both *in vitro* and *in planta*, strains IRDA16, SO16PC and SO16PY were assessed for their bioprotection potential against bacteria canker disease in tomato plants. To evaluate their performance under *in planta* condition, tomato plants were treated with one of the bacterial candidates before (priming) and after (booster) inoculation with *Cm*. Bacterial canker symptoms were observed for all inoculated tomato plants. However, *P. agglomerans* SO16PY delayed the onset of disease by 7 days (18-20 dpi), compared to the *Cm* only-inoculated control (11-13 dpi) (Figs. 4 and 5). For tomato plants pre-treated with *P. marginalis* strains IRDA16 and SO16PC, the first symptoms were observed 15-17 dpi. In general, the severity of disease was lower in tomato plants treated with *P. agglomerans* than with *P. marginalis*. In tomato plants which were treated with *P. agglomerans,* wilting symptoms were observed in only one branch, while for *P. marginalis*, 2-3 branches showed wilting symptoms. For non-treated plants (only *Cm* inoculation), most of leaves and branches showed wilting symptoms (Fig. 4). In plants treated with *P. agglomerans* SO16PY, marginal leaf chlorosis was observed in multiple leaves but no wilting symptoms were detected. Disease severity (DS) score reached 85% for *Cm*-infected plants (without treatment), while this value was 67.5%, 57.5 % and 45% when the plants were treated with *P. marginalis* IRDA16*, P. marginalis* SO16PC, and *P. agglomerans* SO16PY, respectively (Fig. 5). The maximum DS score of 4 was only observed in untreated tomato plants inoculated with *Cm*. Disease incidence was 100% (5/5) for all *Cm*-inoculated plants. Negative control tomato plants inoculated with sterile distilled water (SDW) remained healthy at 28 dpi and no symptoms were observed (Figs. 4 and 5).

**Figure 4.**
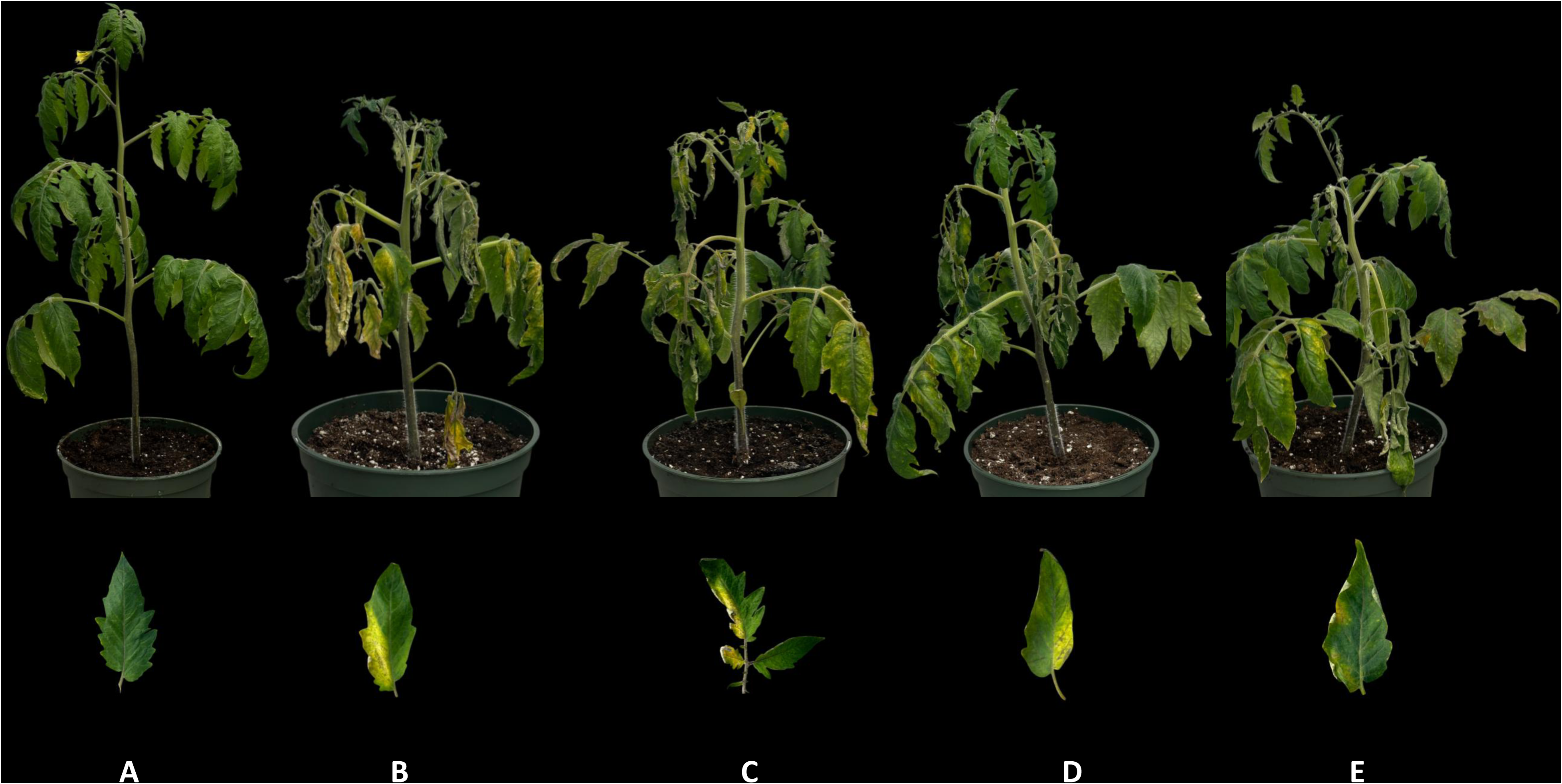
Biocontrol potential of bacterial strains on *Cm* infection in tomato plants. Treatments were as followed: **A.** Negative control (sterile distilled water), **B.** *Cm* inoculation (LMG7333^T^), **C.** *Cm*-IRDA16, **D.** *Cm*-SO16PC, and **E.** *Cm*-SO16PY. Pictures were taken 28 days post-inoculation.

**Figure 5.**
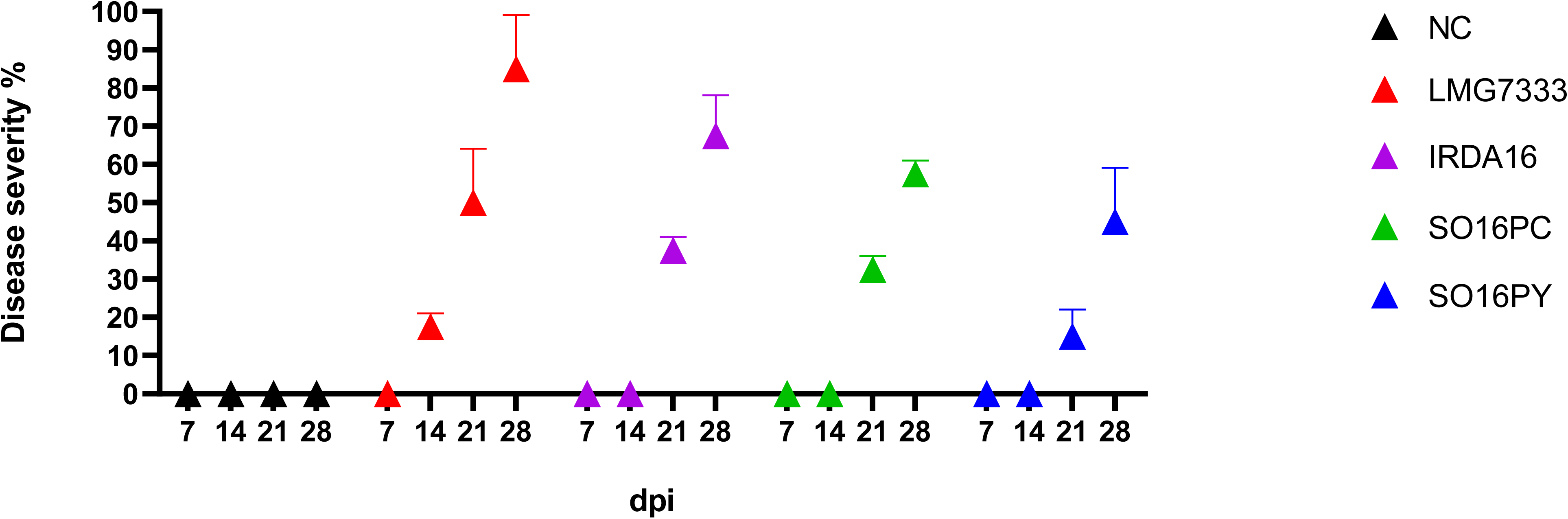
Disease severity of bacterial canker caused by *Cm* after treatment with candidate PGPR. Results up to 28 days post-inoculation. Bars indicate means ± standard errors (n=10), from two replicates. **NC:** negative control (Sterile distilled water).

### Whole genome sequence analyses

To further characterize the selected strains beyond 16s rDNA sequencing and to gain deeper insight into genomic determinants associated with plant growth-promoting traits and secondary metabolite biosynthesis, whole genome sequencing was performed for the selected strains. Genome analysis validated the genus identification obtained by 16S sequencing. Genomic characteristics of the three strains are presented in Table 2. In an ANI-based phylogenetic tree, strain SO16PY clusters with type strain *P. agglomerans* LMG 1286^T^ (ANI: 98.3%; dDDH: 89.1%; Fig. 6A). On the other hand, strains IRDA16 and SO16PC cluster with two strains identified as *P. marginalis* isolated respectively from the rhizosphere of wheat (MGMM3) and sugar beet (ORh26), suggesting their difference from *P. marginalis* sensu stricto (Fig. 6B). Accordingly, ANI (≈96.3%) and DDH indices (≈69.5%, below the threshold level of 70%) were calculated for strains IRDA16, SO16PC, MGMM3 and ORh26 compared with *P. marginalis* type strain DSM 13124^T^. The heatmap based on ANI and DDH (Fig. 7) demonstrates that the observed values are consistent with the hypothesis that these four strains belong to a distinct phylogenic cluster. Our results further highlight heterogenicity in the complex classification of this species (17).

**Figure 6.**
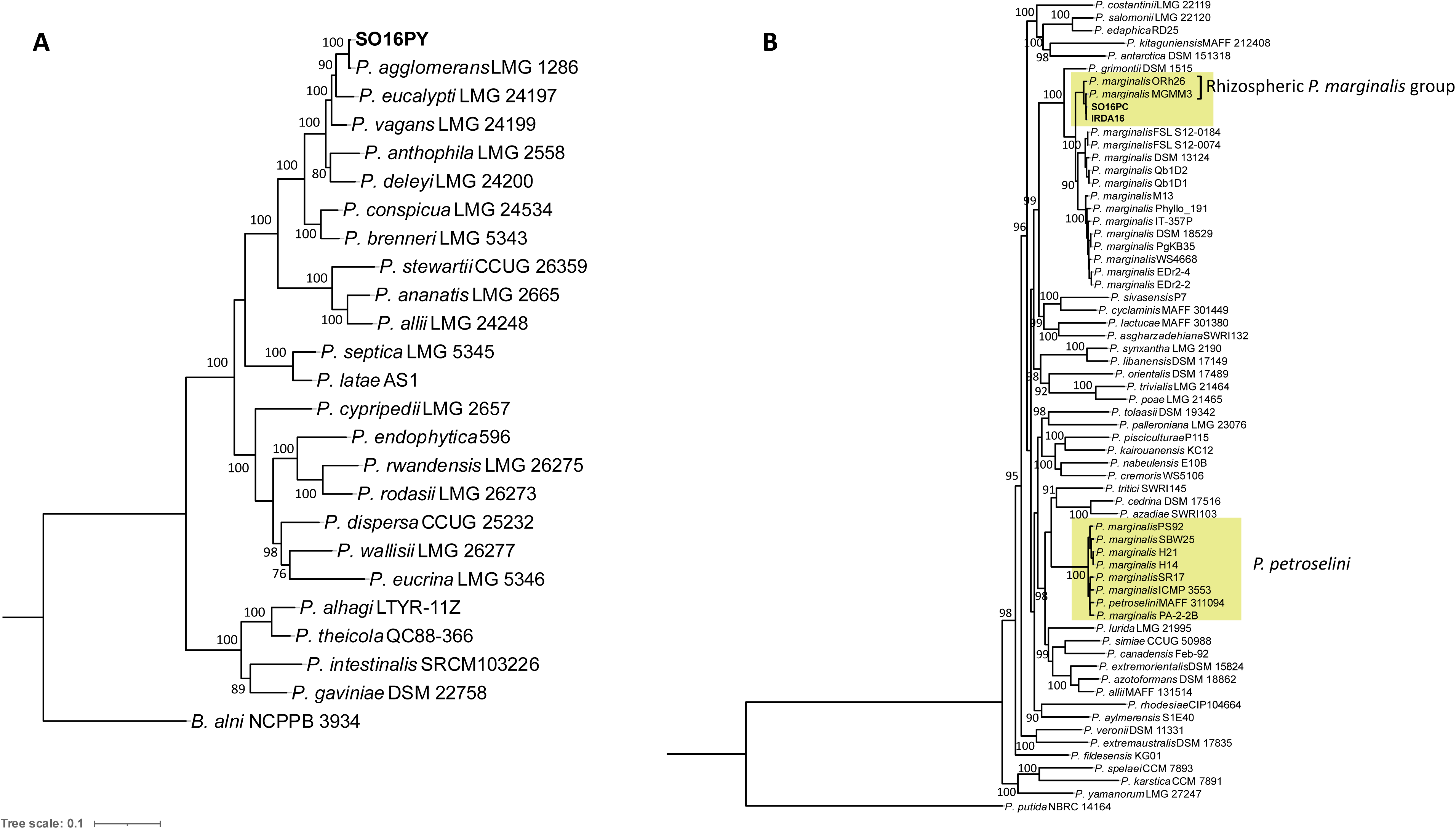
ANI-based maximum likelihood phylogenetic tree of strains SO16PY, IRDA16 and SO16PC. Trees generated using IQ-TREE (1,000 bootstrap replicates). **A.** *Pantoea* spp. and **B.** *Pseudomonas fluorescens* group. Bootstrap values are indicated at each node. *Brenerria alni* NCPPB3934 and *Pseudomonas putida* NBRC14164 were respectively used as outgroup for the *P. agglomerans* and *P. fluorescens* groups.

**Figure 7.**
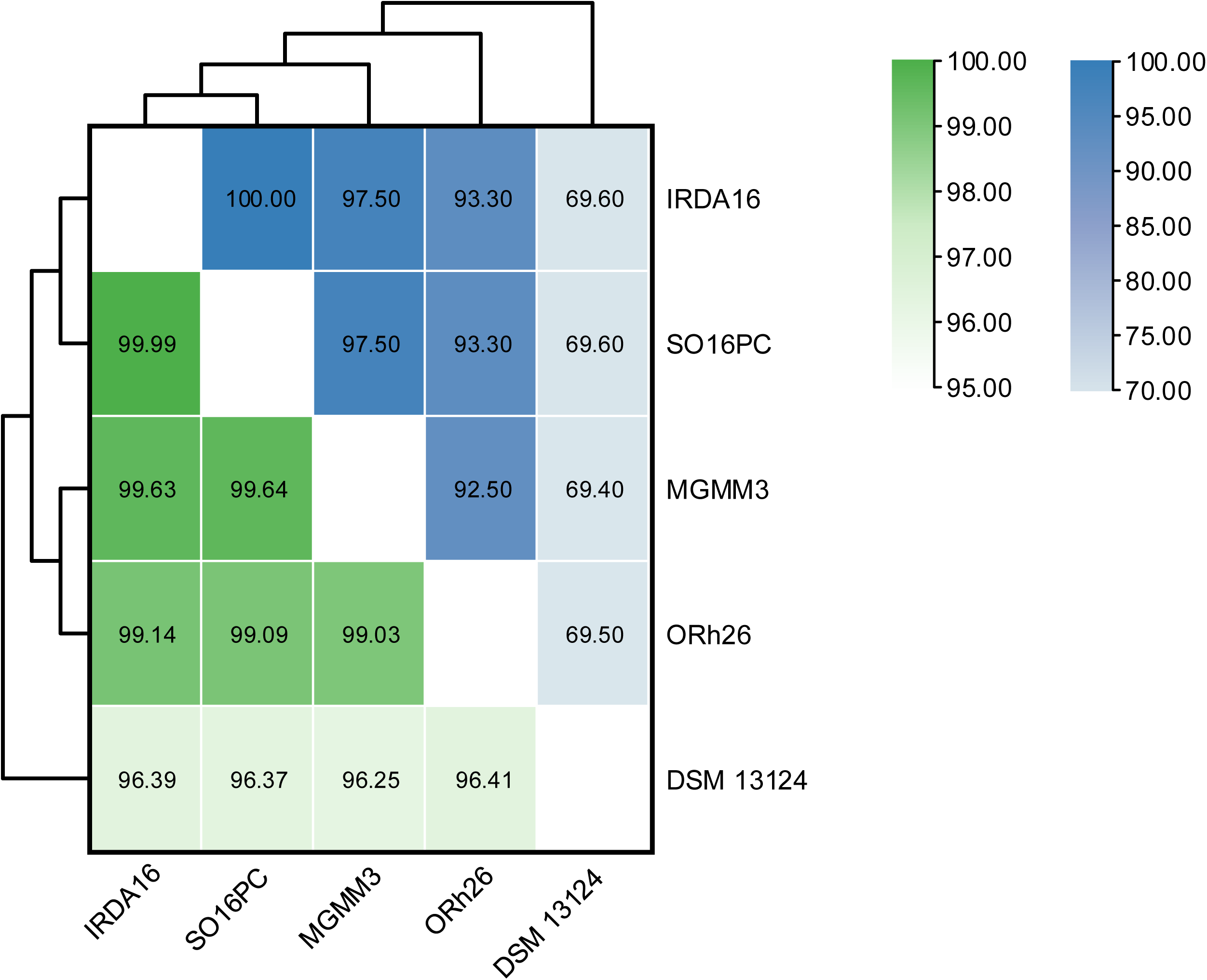
**Heatmap illustrating pairwise genomic similarity based on average nucleotide identity (ANI) and digital DNA-DNA hybridization (dDDH) values**. Studied strains are IRDA16, SO16PC, MGMM3, ORh26 and reference genome DSM 13124^T^. ANI values (%) are indicated in green, while dDDH values (%) are shown in blue. The dendrogram illustrates the hierarchical clustering of strains based on similarity patterns.

**TABLE 2.** Genomic characterization of bacterial strains sequenced in this study*.

| Strain | IRDA16 | SO16PC | SO16PY |
| --- | --- | --- | --- |
| **Genome size (bp)** | 6,683,402 | 6,684,009 | 5,113,447 |
| **GC content (%)** | 61.2 | 61.2 | 54.9 |
| **No. CDSs** | 6104 | 6103 | 4720 |
| **No. tRNA** | 54 | 54 | 59 |
| **No. rRNA** | 4 | 5 | 4 |
| **No. ncRNA** | 4 | 4 | 12 |
| **No. Pseudo Genes** | 81 | 84 | 59 |

\* Genome annotation was performed using the NCBI Prokaryotic Genome Annotation Pipeline (PGAP).

In total, 4720, 6104, and 6103 protein-coding sequences (CDSs) were respectively identified for strains SO16PY, IRDA16, and SO16PC (Table 2). Orthologous protein clusters were determined for each strain using the OrthoVenn online service. Strain *P. agglomerans* SO16PY shares 3955 proteins with reference strain LMG 1286^T^ (Fig. 8A). SO16PY encodes for 36 unique proteins in its genome which were annotated for DNA recombination, RNA binding, SOS response, sorbitol catabolic process and some unknown functional clusters. Interestingly, strains IRDA16 and SO16PC share 397 proteins with strains MGMM3 and ORh26 that are not found in DSM 13124^T^, while the latter has 45 CDSs that are unique to this strain (Fig. 8B). The protein orthologous groups assignment is presented in Figure 9.

**Figure 8.**
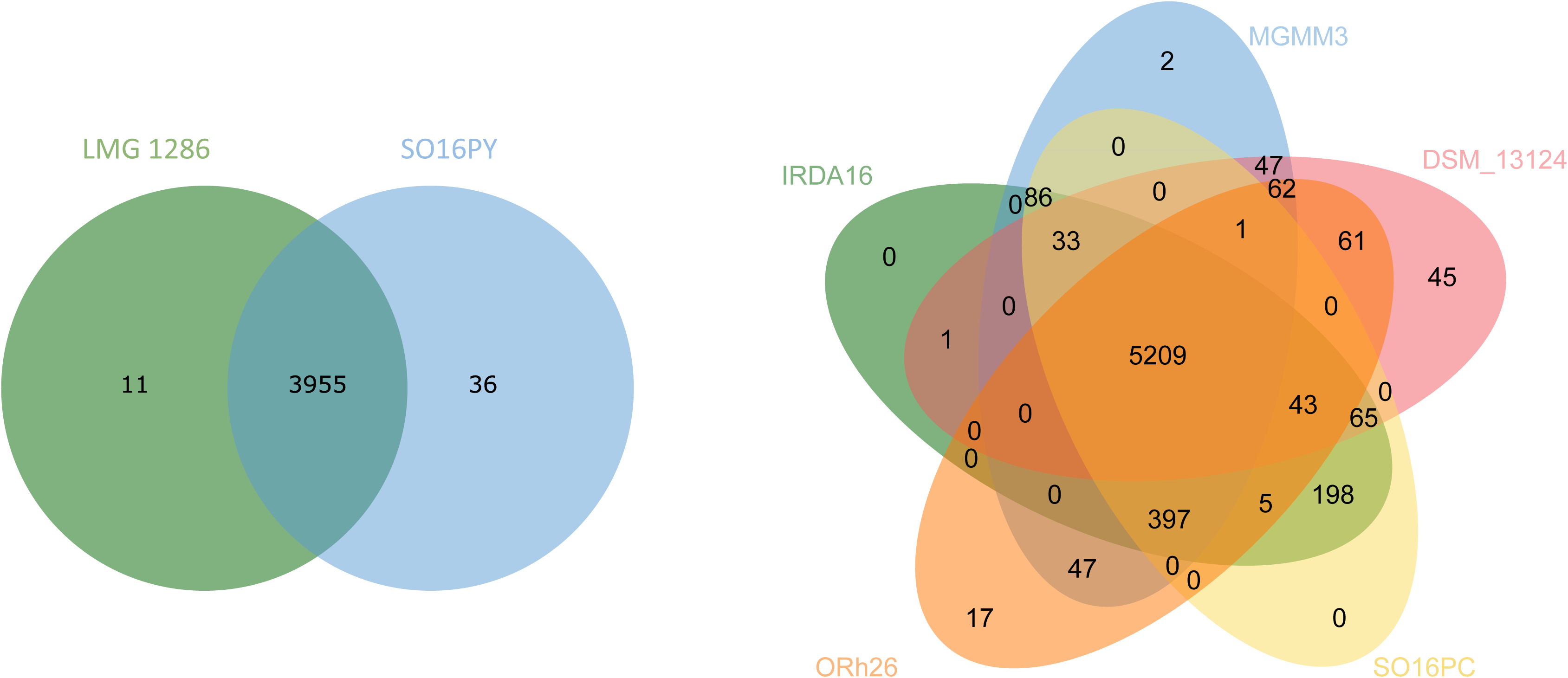
Venn diagrams showing the distribution of shared gene families. Constructed using the OrthoVenn online service. **A.** *Pantoea agglomerans,* and **B.** *Pseudomonas marginalis*. Numbers represent protein numbers, which are common or unique to each strain.

**Figure 9.**
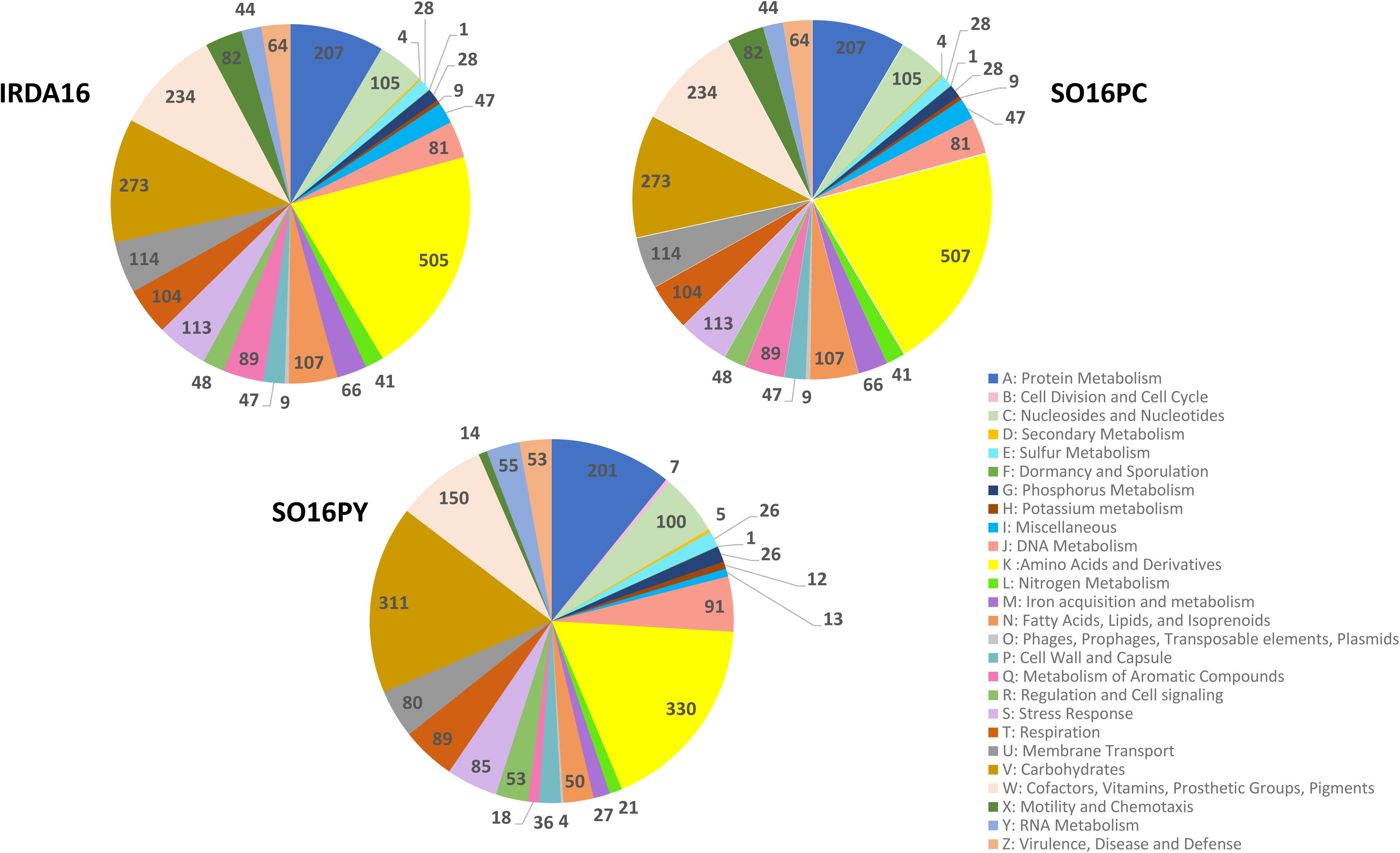
Clusters of orthologous groups (COG) of the annotated CDSs of bacterial strains. The SEED-Viewer version 2.0 was used for the identification and prediction of gene clusters in the genomes. Annotated genes were classified into 26 categories. The numbers next to each section indicate the number of genes in each COG cluster.

### Identification of plant-growth promotion functional genes

To investigate the genetic basis of PGP traits in the sequenced bacterial genomes, protein annotation was performed using the PGPT-Pred tool available on the PLaBAse server (18). Identified PGP-related proteins were further validated and compared with PGAP and Bakta annotation outputs. Genes involved in siderophore production, such as non-ribosomal peptide synthetases (NRPS), *iucA/iucC* (NRPS-independent siderophore (NIS)), *pvdS* and *pvdP* (pyoverdine biosynthesis proteins), *tonB* (TonB-dependent transporter involved in iron uptake) and *fur* (ferric uptake regulator controlling iron homeostasis) were found in both *P. marginalis* strains (IRDA16 and SO16PC). Genes *dfoA* (desferrioxamine biosynthesis protein A), *dfoC* (desferrioxamine biosynthesis protein C), *fhuA-F* (ferric hydroxamate uptake proteins), *fur* and *tonB* were detected in the genome of strain SO16PY (Fig. 10). These results revealed that siderophore production in *Pseudomonas* strains (IRDA16 and SO16PC) is associated with NRPS-dependent (e.g., pyoverdine) and NRPS-independent (e.g., hydroxamate-type) pathways, whereas *Pantoea* (SO16PY) relies on a desferrioxamine-related (e.g., hydroxamate-type) pathway. All genes and their corresponding functions are listed in Table S3.

**Figure 10.**
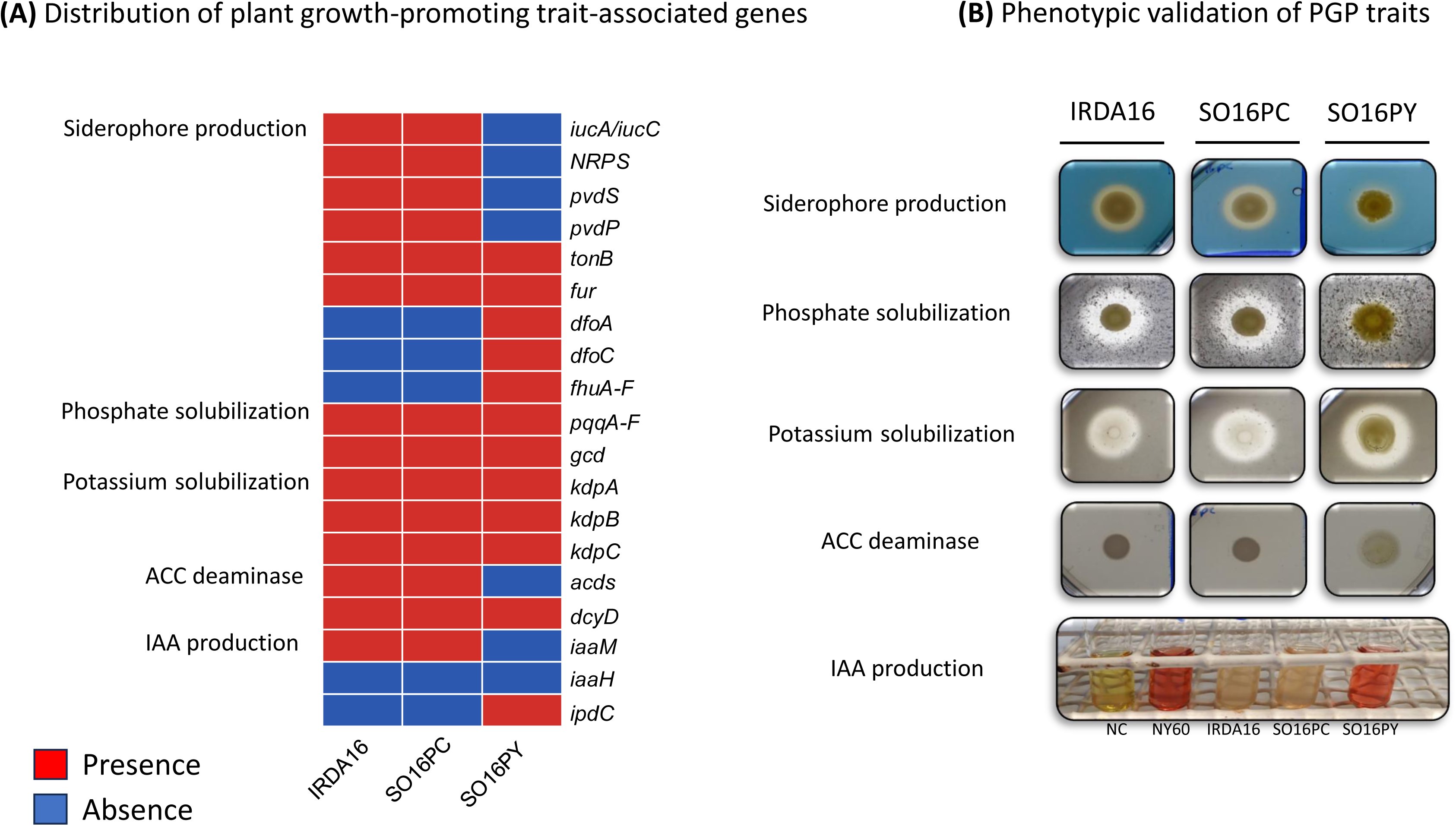
Comparative analysis of plant growth-promoting traits among bacterial strains IRDA16, SO16PC, and SO16PY. **A.** Distribution of genes associated with siderophore production, phosphate and potassium solubilization, ACC deaminase activity, and indole-3-acetic acid (IAA) production. Red indicates gene presence and blue indicates absence. **B.** Phenotypic validation of corresponding PGP traits. Detailed information on the identified genes is provided in Table S3.

All three strains display phosphate and potassium solubilization activities (Table 1, Fig. 10). Genes involved in phosphate solubilization *i.e*. *pqqA-F,* (pyrroloquinoline quinone biosynthesis proteins), and *gcd* (glucose dehydrogenase) were found in all three strains. The *pqqA–F* gene cluster is responsible for the biosynthesis of the redox cofactor PQQ, which is required for the activity of *gcd*, an enzyme that catalyzes the oxidation of glucose to gluconic acid, thereby contributing to phosphate solubilization.

PGPT-Pred also identified *acdS* (1-aminocyclopropane-1-carboxylate deaminase) in *P. marginalis* and *dcyD* (D-cysteine desulfhydrase), which is a homolog of ACC deaminase genes, in *P. agglomerans* (19). The presence of *acdS*, encoding ACC deaminase, suggests the potential to lower ethylene levels through ACC degradation, thereby promoting plant growth.

For production of the auxin IAA, our results indicate that strain SO16PY uses the IAA pathway *i.e.* through indole-3-pyruvic acid (IPyA). In this pathway, tryptophan is first transaminated to indole-3-pyruvic acid, which is then decarboxylated to indole-3-acetaldehyde, followed by oxidation to IAA. Genes like *ipdC* (Indole-3-pyruvate decarboxylase) are critical in this pathway, which was detected on the genome. For strains IRDA16 and SO16PC, the indole-3-acetamide (IAM) pathway (*iaaM* and *iaaH*) was identified. This pathway involves the conversion of tryptophan to indole-3-acetamide by tryptophan monooxygenase (encoded by *iaaM*) and then to IAA by indole-3-acetamide hydrolase (product of *iaaH*). However, in our *Pseudomonas* genomes only *iaaM* was detected and accordingly no IAA production activity was observed *in vitro* (Table 1 and Fig. 10).

### Identification of biosynthetic gene clusters

To further explore the biosynthetic potential of the selected bacterial strains, whole-genome sequences were analyzed using the antiSMASH database to identify secondary metabolite biosynthetic gene clusters (BGCs) (20). Particular attention was given to clusters associated with the production of antimicrobial compounds, siderophores, and other bioactive metabolites that may contribute to PGP and biocontrol activity. This analysis provided additional insights into the functional capabilities of the isolates and their potential ecological roles. The summary of the predicted secondary metabolites for each strain is presented in Table S4. In *P. agglomerans* SO16PY, genes responsible for the biosynthesis of desferrioxamine E, frederiksenibactin (a metallophore), an aryl polyene pigment and phenazines were detected. In *P. marginalis* IRDA16 and SO16PC, genes that could be involved in the production of several unknown NRPS were revealed by the analysis. In addition, genes for the lipopeptide viscosin and a terpene such as sodorifen, were also detected (Table S4).

## DISCUSSION

Biocontrol approaches are considered an attractive alternative over chemical pesticides since they have lower toxic effects on non-target organisms and can reduce negative impact on biodiversity (21). Beneficial microorganisms, particularly PGPR, have therefore emerged as important tools in biocontrol and sustainable crop protection strategies. These bacteria enhance plant growth and contribute to the suppression of phytopathogens (3,5). In this study, we aimed to identify bacterial strains with combined PGP and biocontrol activity against the phytopathogen *Cm,* the casual agent of bacterial canker disease on tomato plants. Following *in vitro* and *in planta* screenings, two strains of *Pseudomonas marginalis* (IRDA16 and SO16PC) and one strain of *Pantoea agglomerans* (SO16PY) showed potential to promote tomato growth and control *Cm*. This stepwise approach allowed us to identify a limited number of effective bacterial candidates with consistent performance under controlled growth chamber conditions, highlighting the importance of *in planta* validation for functional screening of beneficial microorganisms and indicating that *in vitro* screening alone is not sufficient to predict *in planta* performance due to environmental complexity and plant–microbe–soil interactions (22,23).

Our results from *in planta* test showed that *P. agglomerans* SO16PY is able to improve tomato growth parameters for both tested cultivars, Trust and Heinz 2206, under less rich soil condition (PROMIX: sand, 70:30, v/v). Soltani et al. (2025) previously reported that a *P. agglomerans* isolate enhanced tomato growth and yield, as well as nutritional properties (24). Likewise, Vasseur-Coronado et al. (2021) demonstrated that volatile organic compounds emitted by a *P. agglomerans* increased lateral root density, root and shoot dry weight of tomato seedlings (25). Consistent with these findings, improvements in shoot length and shoot dry weight were also reported in tomato plants treated by another *P. agglomerans* (26). Beyond its PGP effects, our results from *in planta* biocontrol test show that *P. agglomerans* SO16PY not only can delay onset of *Cm* disease progression in tomato plants but also decrease bacterial canker disease severity (wilting). Although, several leaves showed marginal leaf chlorosis, the severity of wilting was much lower compared to control plants inoculated with *Cm*. Our findings are consistent with previous reports showing that *P. agglomerans* can reduce by 70% the onset of bacterial canker disease symptoms on tomato plants (27,28). Together with its strong inhibitory effect against *Cm in vitro,* compared to other strains (Table S1), these findings suggest that *P. agglomerans* could be a promising PGPR candidate for controlling *Cm* in tomato plants, particularly under less rich soil conditions, in line with multiple studies that have highlighted both the biocontrol and biostimulant potential of *P. agglomerans* isolates (24,25,29,30).

Two *Pseudomonas marginalis* strains, IRDA16 and SO16PC, exhibited PGP activity on tomato plants, although their effects varied depending on cultivar and soil conditions. Both strains enhanced root and shoot biomass in the cultivar Trust under standard PROMIX condition, indicating their capacity to promote plant growth under nutrient-rich substrates. In contrast, strain IRDA16 showed a more cultivar-specific effect, improving growth parameters primarily on Heinz 2206, while SO16PC maintained its positive impact on shoot biomass even under less rich soil condition (PROMIX: sand, 70:30, v/v). This suggests that SO16PC may possess greater adaptability to suboptimal growth environments, an important trait for field applications where soil quality is often limiting.

Despite promising antagonistic activity *in vitro*, the biocontrol performance of IRDA16 and SO16PC against bacterial canker was less pronounced than *P. agglomerans* SO16PY. Both *P. marginalis* strains delayed symptom onset by several days and reduced disease severity relative to the untreated control (Fig. 5), confirming their capacity to interfere with disease progression. However, the extent of protection remained limited, as multiple branches still exhibited wilting symptoms and overall disease severity scores were higher than those observed for *P. agglomerans*-treated plants. Taken together, this information suggests that while *P. marginalis* strains contribute to disease suppression, their mechanisms of action or efficiency may differ from or be less pronounced than those of SO16PY. It also suggests that IRDA16 and SO16PC may be more effective as part of integrated strategies rather than as standalone inoculants.

Interestingly, strain SO16PC which was isolated from tomato showed better potential than strain IRDA16 (isolated from apple leaf) to reduce the wilting symptoms caused by *Cm* (Fig. 5). This observation suggests that the original host of isolation may influence the adaptability and functional performance of bacterial strains in plant–microbe interaction. Interestingly, *P*. *marginalis* strain ORh26 (isolated from rhizosphere of sugar beet), which clustered within the same newly identified rhizosphere-associated lineage as strains IRDA16 and SO16PC in our phylogenomic analysis (Fig. 6B), was recently reported to induce systemic resistance in sugar beet and reduce pathogen proliferation and leaf lesion size caused by *Pseudomonas syringae* pv. *aptata* (31). Consistent with these findings, biocontrol and PGP activities of *P. marginalis* isolates have also been documented in several studies (32,33,34,35). For instance, a *P. marginalis* isolate (from mixture of peat and compost) was able to reduce the severity of the tomato root rot disease caused by *Pythium ultimum* in an hydroponic system while also improving growth parameters and fruit production of infected tomato plants compared to control inoculated with *P. ultimum* (36).

To clarify the phylogenetic relationships and assess the taxonomic status of the studied strains, whole-genome sequencing analysis was conducted. Based on ANI and digital DNA–DNA hybridization (dDDH) indices, our *Pseudomonas* strains showed the highest similarity to members of the *P. marginalis* within *Pseudomonas fluorescens* subgroup (*P. fluorescens* group), with values close to the species delineation thresholds (ANI≈96% and dDDH≈69.5%). Therefore, complete genome sequences of all representative strains from this group were included in the phylogenetic analysis (Fig. 6B) to clarify their affiliation (37,38). Phylogenomic analyses revealed that our two strains form a new distinct lineage together with the rhizosphere-associated strains *P. marginalis* MGMM3 and ORh26, clearly separated from the classical pathogenic lineage represented by the type strain and other disease-associated *P. marginalis* isolates. Although ANI values (≈96%) placed these strains within the *P. marginalis* species boundary, the borderline dDDH value (≈69.5%), consistent separation in the phylogenetic tree, and presence of 397 unique protein clusters suggest the emergence of a specialized rhizosphere lineage with a lifestyle different from the pathogenic type strain. Unlike the canonical pathogenic lineage, which is primarily associated with plant tissue invasion and disease development, these strains exhibited PGPR-associated traits and genomic features linked to beneficial plant–microbe interactions. Based on this phylogenomic analysis, we propose that strains IRDA16, SO16PC, MGMM3, and ORh26 represent a novel *Pseudomonas* species, following the recommendation of Nedeljković et al. (2025) to reclassify strain ORh26 (31). It is probable that this population is undergoing genetic divergence. Recent phylogenetic analysis performed by Sawada et al. (2023) confirmed that *P. marginalis* sensu lato is heterogeneous, suggesting that it might be a species complex containing many cryptic species (17). In phylogenetic tree (Fig. 6B), several bacterial strains (ICMP 3553, PS92, SBW25, H21, H14, SR17, MAFF 311094 and PA-2-2B) previously identified as *P. marginalis* in the NCBI database, clustered with *P. petroselini,* the causal agent of bacterial rot of parsley and spinach. Heatmap illustrating pairwise genomic similarity based on average ANI and dDDH values confirmed the relationships inferred from the phylogenetic tree (Fig. 11). This observation aligns with the findings of Sawada et al. (2023), who proposed the reclassification of strains ICMP 3553,

**Figure 11.**
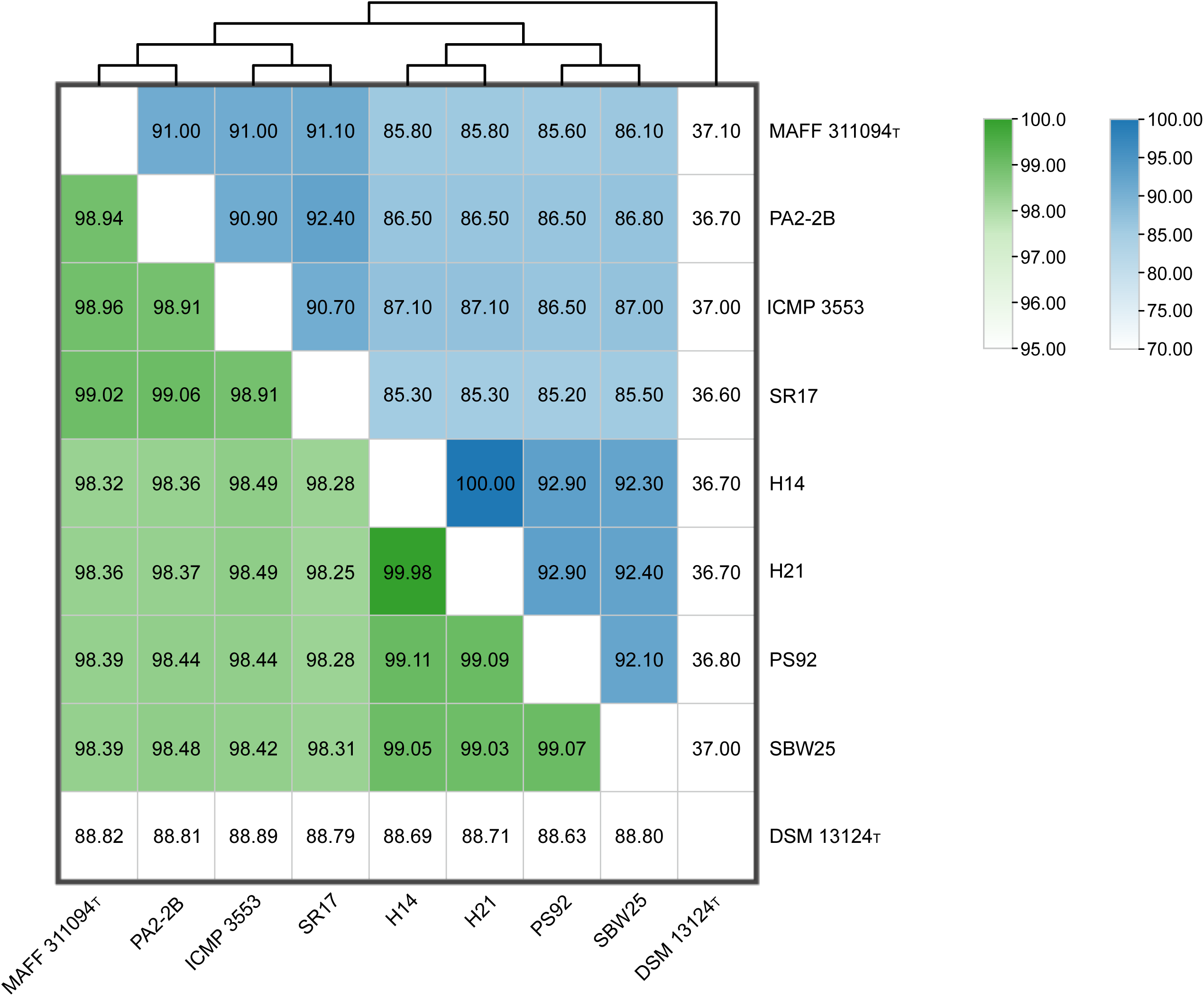
Heatmap illustrating pairwise genomic similarity based on average nucleotide identity (ANI) and digital DNA-DNA hybridization (dDDH) values among the studied strains. IRDA16, SO16PC, MGMM3 and ORh26 are compared to reference genomes *Pseudomonas marginalis* DSM 13124^T^ and *Pseudomonas petroselini* MAFF 311094^T^. ANI value (%) are indicated in green, while dDDH values (%) are shown in blue. The dendrogram illustrates the hierarchical clustering of strains based on similarity patterns.

MAFF 311094, and SR17 as *P. petroselini*. A PGPR strain *P. marginalis* SBW25, historically classified as *Pseudomonas fluorescens* SBW25, has been documented for its PGP and biocontrol activity (39,40,41,42). The placement of this strain within the *P. petroselini*-related clade further supports the presence of non-pathogenic, plant-beneficial lineages within taxa traditionally associated with disease. However, a formal taxonomic assessment of this group was beyond the scope of our study, and additional research is suggested to clarify the taxonomic status of this complex lineage (17,43).

Based on whole genome sequence analysis, we found several BGCs in sequenced bacterial genomes for the production of an array of bioactive metabolites. Since the cell free supernatants of the three bacteria were effective in inhibiting the growth of *Cm*, it can be inferred that these compounds might be involved in biocontrol activity. For instance, in *P. agglomerans* we identified the desferrioxamine E BGC, a siderophore of the hydroxamate group (44). Choi et al. (2022) confirmed that this siderophore is required for antibacterial activity of *Pantoea ananatis* against *Erwinia amylovora* (45). Phenazine production genes were also found in the genome of strain SO16PY. Phenazines are redox-active molecules that display broad-spectrum antibiotic activity toward many bacterial, fungal, and oomycete plant pathogens (46).

We found several predicted BGCs in the *P. marginalis* sequenced genomes coding for unknown metabolites; it remains to be determined which ones are involved in antagonistic activity against *Cm*. Genes responsible for the lipopeptide viscosin were identified, showing high similarity to those found in *P. marginalis* strain SBW25 (Table S4). Several roles have been associated with viscosin production, including motility, root colonization and PGP activity (39,40). Interestingly, none of the biosynthetic gene clusters involved in production of viscosin, terpenes, thanamycin, nunapeptin, or crochelin A (Table S4) are present in the genome of type strain DSM 13124^T^ (Table S5). Given the well-established role of viscosin in promoting plant growth and terpene production in plant defence, the acquisition of these biosynthetic genes may have been a key factor in the ecological adaptation of strains IRDA16 and SO16PC. In particular, viscosin production could enhance biocontrol capabilities while also improving rhizosphere competence through increased motility, biofilm formation, and more efficient root colonization, ultimately supporting a beneficial plant-associated lifestyle (39,47,48). A conceptual model summarizing these relationships is presented in Figure 12.

**Figure 12.**
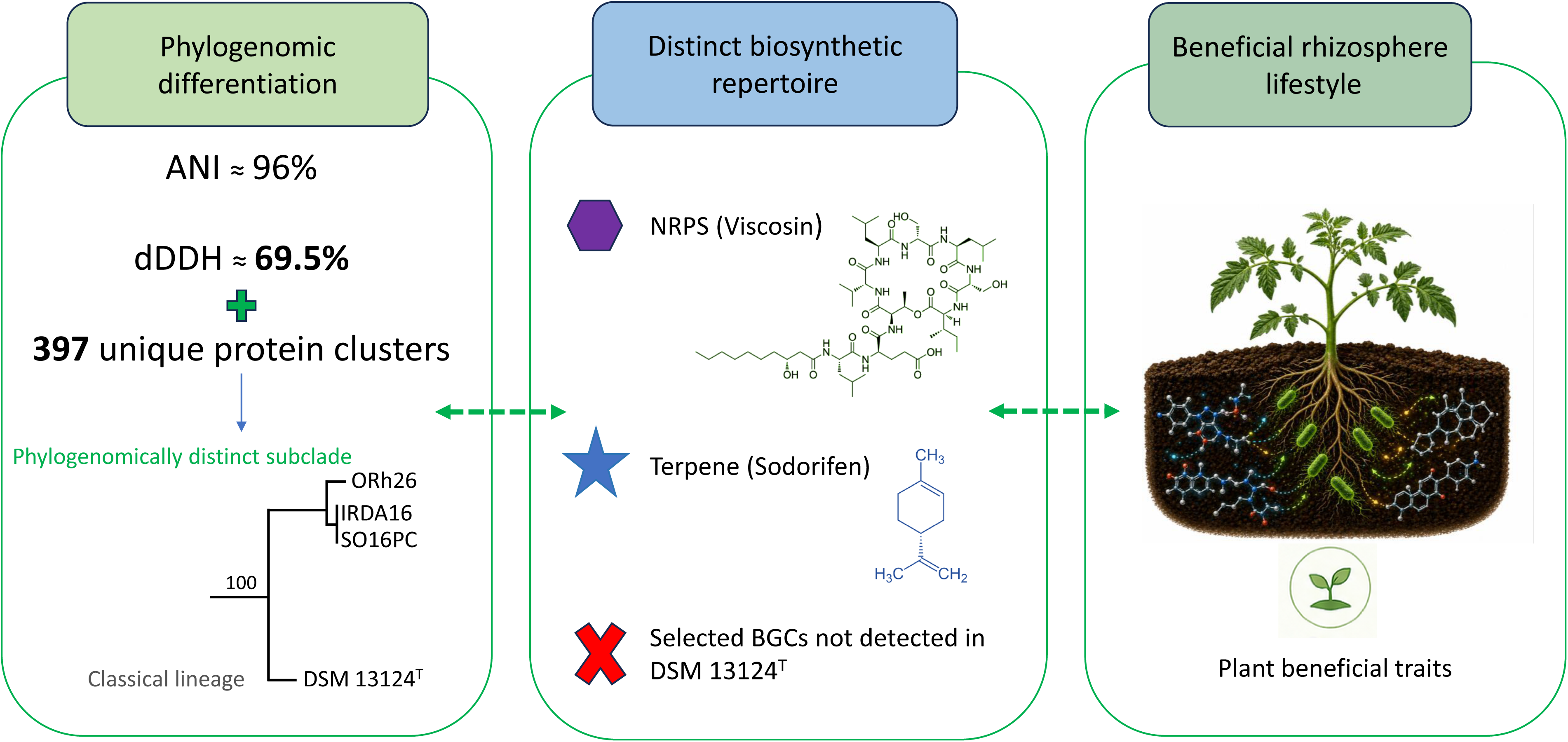
A conceptual model integrating phylogenomic divergence and plant beneficial phenotypes. The model illustrates a proposed (hypothesized) association between genomic differentiation, the presence of unique biosynthetic gene clusters (BGCs), and beneficial rhizosphere lifestyle observed in the studied strains.

To date, a few bacterial strains have been reported to control *Cm* and promote tomato growth as well (3). For instance, a *Pseudomonas putida* isolate applied as a seed and root treatment significantly reduced the severity of bacterial canker in tomato plants (49). Similarly, strains of *Bacillus* spp. and *Serratia* sp. exhibited an antagonistic activity against *Cm in vitro* and significantly reduced the incidence of tomato bacterial canker in the greenhouse (50,51). In another study, Gautam et al. (2019) confirmed the efficiency of *Bacillus amyloliquefaciens* as a biocontrol agent against *Cm* both *in vitro* and *in planta* (52). Our results demonstrate that isolates from other bacterial species can also delay disease development and reduce the severity of wilting symptoms caused by *Cm.* Taken together, the available literature and our findings demonstrate diverse PGPR possess the dual ability to enhance tomato growth and suppress *Cm*, highlighting their potential as sustainable tools for managing this challenging disease. Future research could focus on developing effective microbial consortia and evaluating their performance under diverse environmental conditions and agricultural systems to optimize their reliability and field applicability. In addition, further studies are required to elucidate the mechanisms at play. Work is also necessary to identify the best application method yielding the expected benefits.

## MATERIALS AND METHODS

### Bacterial strains and culture conditions

Bacterial strains from our laboratory collection were used in this study. During 2011-2016, several samplings were done from apple and pear orchards (soil, flower, stem, leaf and fruit), tomato and pepper (soil, leaf and fruit), eggplant (soil), onion (soil), maize (soil), strawberry (soil, leaf and fruit) and raspberry (soil) in Canada (Quebec) and USA (Florida and New York) (53,54,55,56). Routinely, strains were grown in Tryptic soy broth (TSB; Difco) on a rotary shaker (30°C, 200 rpm, 20h), unless otherwise indicated. For *Cm*, type strain LMG7333^T^ (ED4529) was used in all *Cm*-related experiments. For culture preparation, *Cm* cells were grown in TSB medium (30°C, 200 rpm, 30-35h).

### Antagonist activity

Bacterial strains were grown in 5 ml TSB, overnight at 30°C. Then, 5 µl of each culture was spotted on Tryptic Soy Agar (TSA) plates uniformly swabbed with a *C. michiganensis* LMG7333^T^ suspension adjusted to 10^8^ CFU/ml using sterile cotton swabs. The plates were incubated at 30°C for 2 days. The diameter of the inhibition zone around each colony was measured.

To assess the effect of extracellular metabolites on the growth of *Cm*, bacterial cells were removed by centrifugation at 8,000 x *g* for 5 min. The cell free supernatant was then filtrated (0.22 µm). In a Honeycomb^TM^ 100-well plate, cell free supernatant and blank culture medium (TSB) were mixed in final volume of 200 µl at a ratio of 1:1 (v/v) (3 replicates for each treatment).

Strain LMG7333^T^ was inoculated in each well (final OD_600_= 0.3). *Escherichia coli* DH5α (ED78) was used as a negative control. The plate was incubated in a Bioscreen C instrument (Growth Curves USA) for 30h at 30 °C. The OD_600_ was measured every 6h. This experiment was repeated twice.

### Phosphate and potassium solubilization assays

Phosphate solubilization was evaluated on National Botanical Research Institute’s phosphate (NBRIP) medium, prepared as described (57). For the potassium solubilization assay, Aleksandrow medium (HIMEDIA) was used. Bacteria were grown in TSB medium (30°C, 200 rpm, overnight), and cells were harvested by centrifugation at 8,000 x *g* for 5 min. Then, they were washed three times with NaCl 0.8% and suspended to get a concentration of 10^8^ CFU/ml. Agar plates were inoculated with 5 µl of bacterial suspension. The plates were incubated at 28°C. After 7 days, plates were examined for phosphate/potassium solubilization activity by measuring the diameter of the clearing zone surrounding the colonies. This experiment was repeated twice. *Burkholderia vietnamiensis* strain G4 (ED382) was used as positive control in PGPR tests (Table S1) (58).

### Siderophore detection by Chrome-Azurol S (CAS) assay

Bacterial suspension were prepared as described above and were used to inoculate MM9-CAS agar plate (59). King B-CAS was used to assess siderophore production in strains identified as weak producers on MM9-CAS-agar (60,61). Plates were incubated at 28°C for 4 days. The orange halo around colony indicates siderophore production and the diameter of the halo was measured. This experiment was repeated twice.

### ACC deaminase activity

Minimal DF (Dworkin and Foster) salts media (KH_2_PO_4_ [4.0 g L^−1^]; Na_2_HPO_4_ [6.0 g L^−1^]; MgSO_4_.7H_2_O [0.2 g L^−1^], glucose [2.0 g L^−1^], gluconic acid [2.0 g L^−1^]; citric acid [2.0 g L^−1^] and Bacto Agar [18 g L^−1^] with trace elements: FeSO_4_.7H_2_O [1 mg], H_3_BO_3_ [10 mg], MnSO_4_.H_2_O [11.19 mg], ZnSO_4_.7H_2_O [124.6 mg], CuSO_4_.5H_2_O [78.22 mg], MoO_3_ [10 mg], pH 7.2) supplemented with 3 mM ACC (Sigma) was used in this test (62,63). A bacterial suspension was prepared as described earlier and 5 µL were deposited on agar plates. Plates were incubated at 28°C for 3 days. Colonies growing on the plates were taken as ACC deaminase producers. Strains *B. vietnamiensis* G4 (ED382) and *E. coli* DH5α (ED78) were respectively used as positive and negative controls. This experiment was repeated twice.

### Indoleacetic acid (IAA) production

Production of Indole-3-acetic acid (IAA) was determined as previously described with some modifications (64). Bacterial strains were grown overnight in TSB medium at 30 °C. Then, 100 μl of each culture was used to inoculate 6 mL of fresh liquid LB medium supplemented with 5 mM L-tryptophan. Cultures were incubated at 30°C in an orbital shaker (200 rpm) up to 144 h, depending on IAA production ability of the strain. Non-inoculated liquid LB medium (with L-tryptophan) was used as blank. Strain *Pantoea agglomerans* NY60 (ED4177) (Table S1) was used as positive control (54). Each 48 h, 1.5 mL of bacterial culture was taken and centrifuged for 5 min at 16,500 x *g*. Then, 1 mL of each supernatant was mixed with 2 mL of Salkowski reagent (65) and samples were kept in the dark at room temperature for 30 min. The strains for which a pinkish-red color was observed were determined to be producers of IAA. The results were confirmed by measuring the absorbance at 450 (Indole-3-butyric acid (IBA)), 490 (Indole) and 530 (IAA) nm (Cytation 3 microplate Reader, Biotek) (66). A standard curve using an IAA standard (Sigma) at a range of concentrations between 1-100 μg/mL was used for quantification. This experiment was repeated twice.

### 16s rRNA gene sequencing

Genomic DNA of selected bacterial candidates was isolated using a EasyPure Genomic DNA Kit (Transgen) following the manufacturer’s instructions. The partial sequence of the 16S rRNA gene was amplified using primers 27F (5′-AGAGTTTGATCCTGGCTCAG-3′) and 1492R (5′-GGTTACCTTGTTACGACTT-3′)(67). DNA sequencing was performed by the sequencing platform at the Montreal Clinical Research Institute (IRCM). Strain identification to the genus level was carried out using the GenBank database of NCBI (https://www.ncbi.nlm.nih.gov/) and Ezbiocloud (https://www.ezbiocloud.net/).

### PGP effect on tomato growth parameters

Two tomato cultivars: 1) Trust, and 2) Heinz 2206 with two different soil combinations: 1) PROMIX, and 2) PROMIX combined with sand (PROMIX:sand, 70:30, v/v) were used. Tomato seeds were disinfected with 2% sodium hypochlorite (NaOCl) for 2 min, washed several times with sterile distilled water (SDW) and dried under the Biosafety cabinet flow. Tomato seeds were sown in pot in growth chamber (25°C, RH: 55-65%, 16h light and 8h darkness). Bacterial cells grown overnight in TSB medium at 30°C were harvested by centrifugation (8,000 x *g*, 5 min) and resuspend in sterile distilled water (SDW) to final concentration of 10^8^ CFU/ml. Tomato plants (7 replicates/one seedling per replicate) were inoculated (as soil drench) two times with bacterial suspension (Fig. 13A). An initial inoculation of 10 mL per plant was performed when the first true leave started appearing, followed by a second inoculation of 15 mL one week after the first inoculation. The inoculation was performed around the stem-root junction. Seven days after the second inoculation, growth parameters such as fresh root weight (FRW), fresh shoot weight (FSW), dry root weight (DRW) and dry shoot weight (DSW) were measured. No fertilizer was used during the experiment. Data was analyzed with GraphPad Prism 8 (GraphPad Software, San Diego, CA, USA) using One-way ANOVA (p<0.05). The experiment was repeated twice, independently.

**Figure 13.**
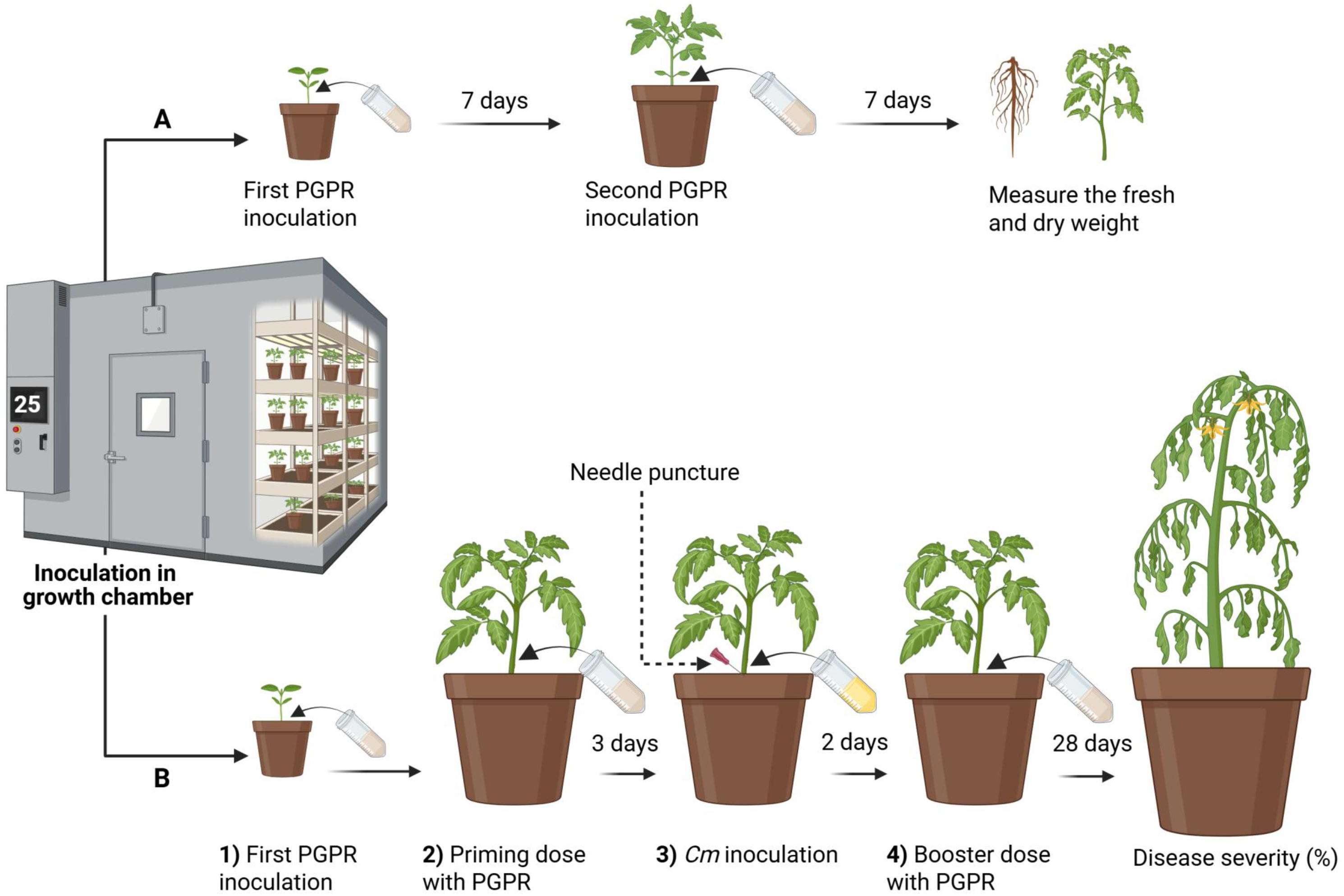
Schematic representation of tomato inoculation methods in growth chamber. **(A)** Plant growth-promoting assay. **(B)** Disease suppression assay.

### PGP effect on tomato seeds

Preparation of tomato seeds and bacterial cells was performed as described above. Tomato seeds were immersed in 10 ml (10^8^ CFU/ml) of each bacterium (gently shaken for 1 h). For negative control, the seeds were soaked in 10 ml SDW. Then, the seeds were dried under the biosafety cabinet flow and placed on freshly wet filter paper inside a Petri dish. A set of each treatment (containing 5 seeds) was replicated two times. The Petri dishes were incubated in a growth chamber at 25°C (10 h light and 14 h darkness) for 7 days. Primary root length was calculated (68,69). Data was analyzed with GraphPad Prism (One-way ANOVA).

### Biocontrol effect to control bacterial canker disease

Tomato seeds (Trust cultivar) were sown in natural soil (PROMIX) and kept in a growth chamber (same conditions mentioned above). The treatments were as followed: 1) Negative control: sterile distilled water (SDW), 2) positive control: *Cm-*inoculated plants, 3) *Cm*+IRDA16, 4) *Cm*+SO16PC, and 5) *Cm*+SO16PY. During the experiment, tomato plants (5 replicates/ strain) were inoculated (as soil drench, 10^8^ CFU/ml) three times with one of the PGPR candidate (Fig. 13B) following these steps: 1) First inoculation was performed when the first true leave start appearing (10 ml/ plant), 2) then, priming dose was added three days before *Cm* inoculation (25 ml/ plant), 3) after that, plants (5 leaf stage) were inoculated as soil drench with *Cm* (5 ml/ plant, 10^8^ CFU/ml). For this, three wounds (needle punctures) were made using a sterile needle around the first stem-root junction and a *Cm* suspension (5 ml/ plant, 10^8^ CFU/ml) was poured in the area where the PGPR treatment had performed, 4) Final booster dose was done two days after *Cm* inoculation (25 ml/ plant). For negative controls, plants were treated with SDW, with the same volume of each treatment for each inoculation point. The progression of the disease was monitored each week, and disease severity (DS) was calculated 28 days post inoculation (28 dpi). Twenty-eight days after inoculation, tomato plants were classified, based on disease symptoms, according to the following scale: 0 ( no symptom); 1 (1-25% leaf chlorosis/wilting); 2 (26-50% leaf chlorosis/wilting); 3 (51-75% wilting); 4 (76-100% of wilting/damping off/seedling death) (70). DS measurements were converted from original ordinal scale to a ratio scale and normalized to 0-1 using the following equation:

DS=[(0×a) +(1×b) +(2×c) +(3×d) +(4×e)]/[(a+b+c+d+e) ×4]

where a, b, c, d and e are the corresponding numbers of infected plants scored from 0 to 4 (71). This experiment was repeated twice, independently.

### Whole genome sequencing

Genome sequencing of strains was performed using Illumina NovaSeq 6000 technologies (SeqCenter, PA, USA). Quality control and downstream processing of the raw reads was performed using the Galaxy platform (http://usegalaxy.org). Trimming and Quality control (QC) were performed using Trimmomatic v0.39 to remove adapter sequences and low-quality reads (72). *De novo* assembly of the filtered reads was performed using SPAdes v4.2 (73). Quality assessment of the assembly was performed using BUSCO v5.8 (74). Based on the assembled sequences, gene prediction and annotation were conducted by Bakta v.1.9.4 (75). Average nucleotide identity (ANI) was calculated among type strains of all species within the genus. The ANI was estimated using OrthoANI online service (76). The dDDH values was estimated using genome-to-genome distances calculator online service (http://ggdc.dsmz.de/distcalc2.php)(77). The accepted threshold for prokaryotic species description, i.e. ≥95 % and ≥70 % for ANI and dDDH, respectively, was used for analysis (78,79,80). Heatmaps illustrating pairwise genomic similarity based on ANI and dDDH values were generated using TBtools (81).

Phylogeny was achieved with Roary v3.13.0 using 80% of identity and IQ-TREE v2.4.0 with bootstrapping (1000 replications) (82,83). The dendrogram was visualized by iTOL v7.1 (https://itol.embl.de/itol.cgi). All bacterial strains used in phylogenetic analysis and their accession numbers are presented in Table S6. Comparisons orthologous clusters were performed using the online service OrthoVenn (http://www.bioinfogenome.net/OrthoVenn/bacteria) (84).

The SEED-Viewer version 2.0 was used for the identification and prediction of represented gene clusters in the genomes (https://rast.nmpdr.org/seedviewer.cgi) (85). Biosynthetic gene clusters (BGCs) were identified using antiSMASH (strict mode) database (https://antismash.secondarymetabolites.org/). PGPT-Pred tool from the PLaBAse server (https://plabase.cs.uni-tuebingen.de/) was used to annotate the plant-growth promoting traits using blastp+hmmer. The results were further verified with PGAP and Bakta annotations and presence of PGP genes was assessed.

## Data availability

The results of whole genome sequences were deposited in the GenBank database of National Center for Biotechnology Information (NCBI https://www.ncbi.nlm.nih.gov/) under accession numbers JBWHZE000000000, JBWJNF000000000 and JBWJNG000000000 corresponding to strains SO16PY, IRDA16 and SO16PC respectively.

## SUPPLEMENTAL MATERIAL

**SUPPLEMENTAL FILES**, Figures S1, Tables S1, S2, S3, S4, S5 and S6.

## ACKNOWLEDGMENTS

This study was funded by the CRIBIQ (grant no. 2022-031-C98) and the Natural Sciences and Engineering Research Council of Canada (Alliance, grant no. ALLRP 583416-23), in partnership with Agro-100 Ltée.

NS and ED conceived and designed the study. NS carried out the experiments. NS analyzed and interpreted the data with assistance from MCG and ED. NS prepared the manuscript. MCG and ED revised the manuscript. All the co-authors contributed to the article and approved the final version of manuscript.

